## Supplemental Materials and Methods for "The Androgen Receptor: A Therapeutic Target in Desmoplastic Small Round Cell Sarcoma"

### Supplemental Figure 1

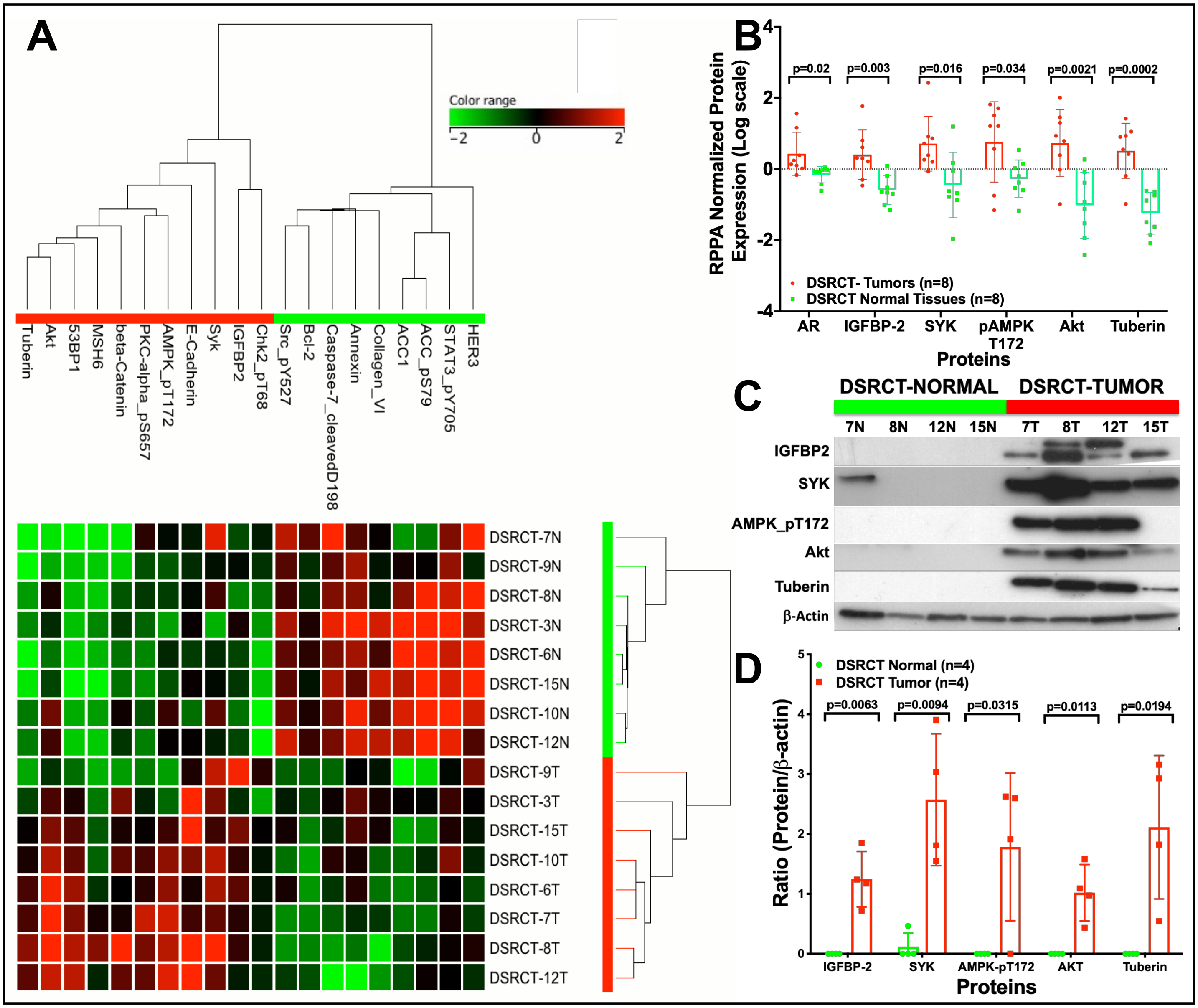

### Supplemental Figure 2

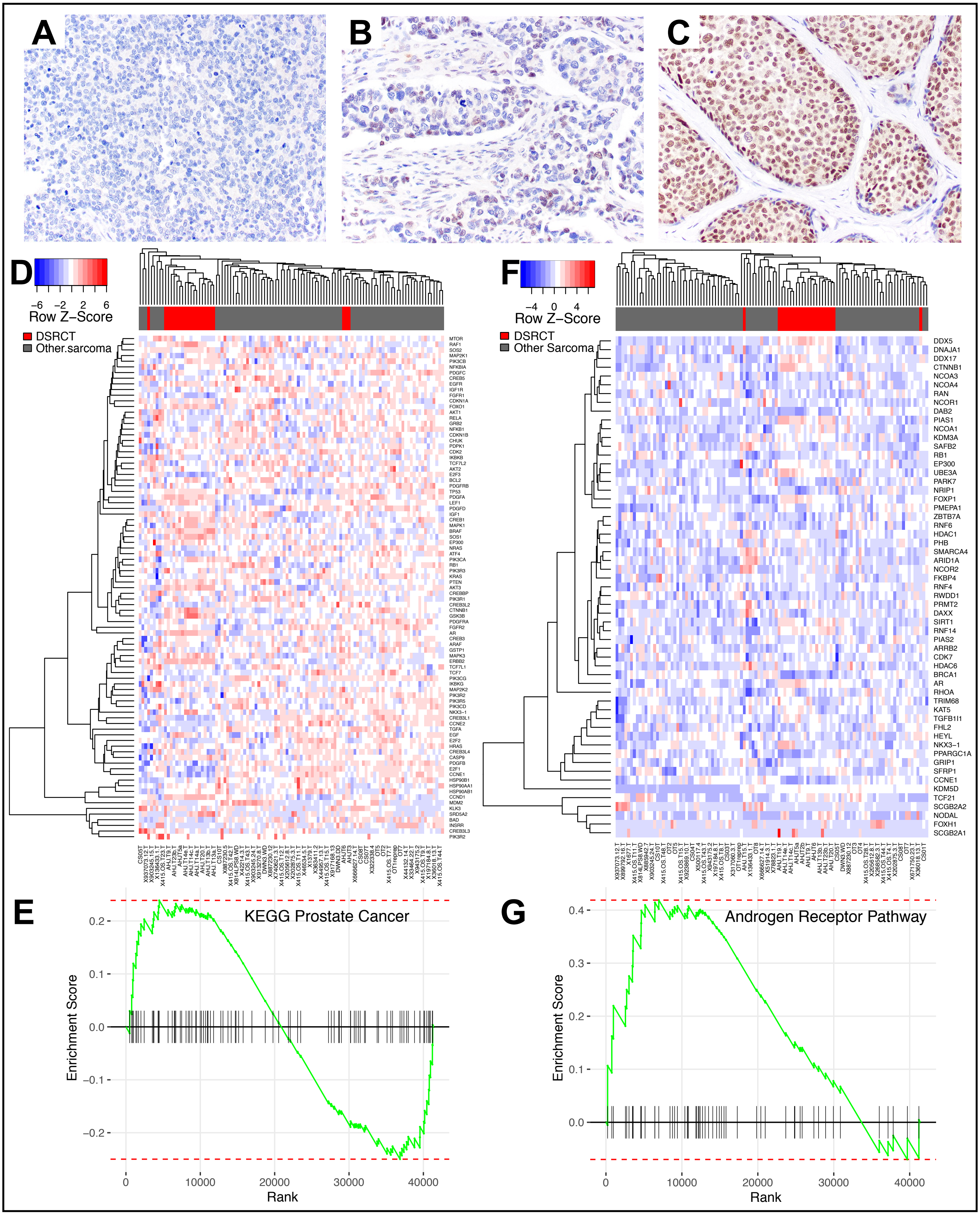

### Supplemental Figure 3

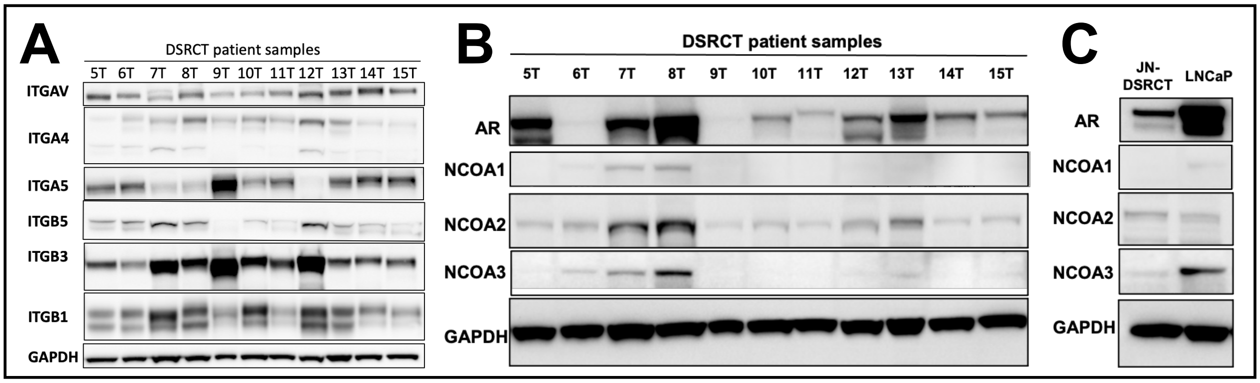

#### Supplemental Figure 4

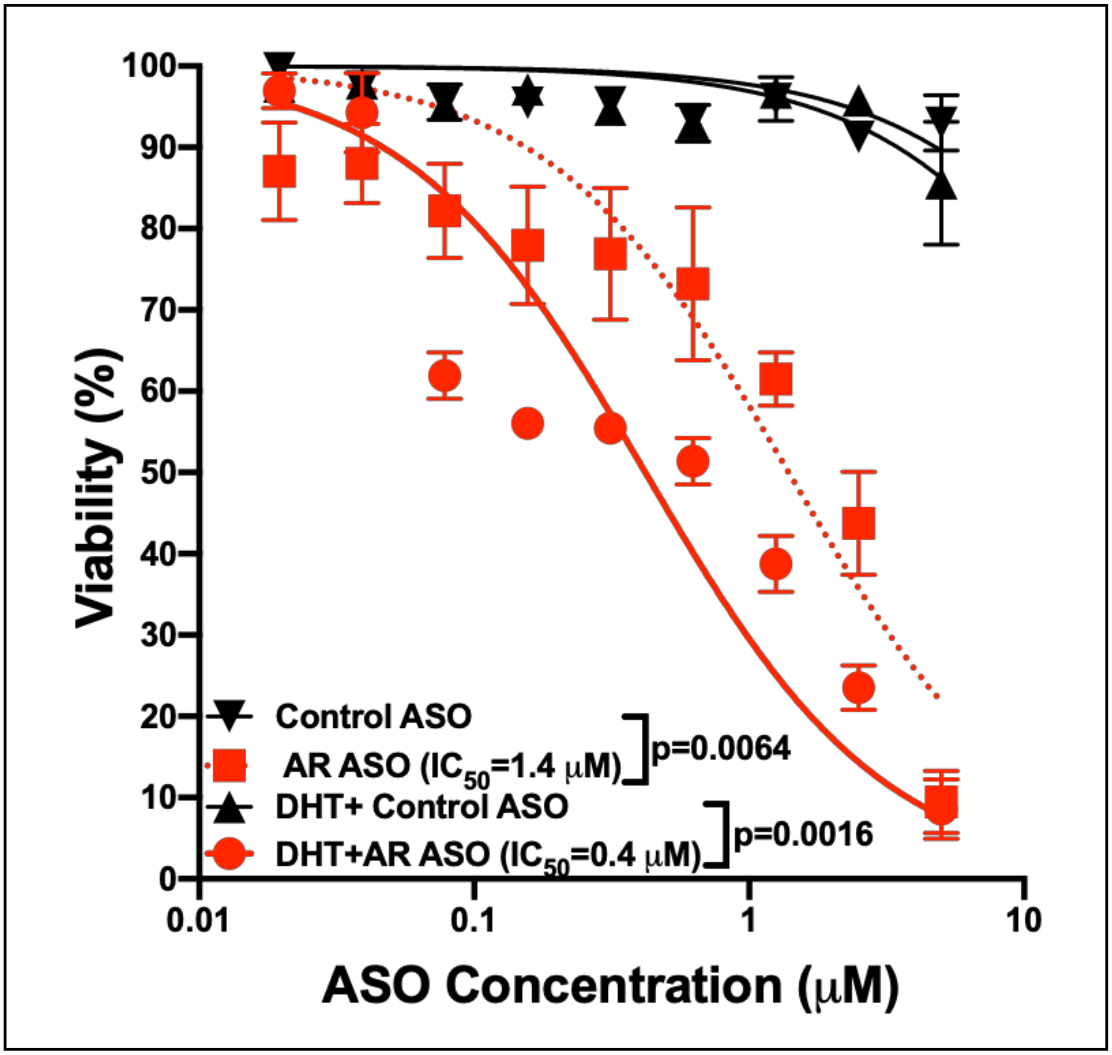

### Supplemental Figure 5

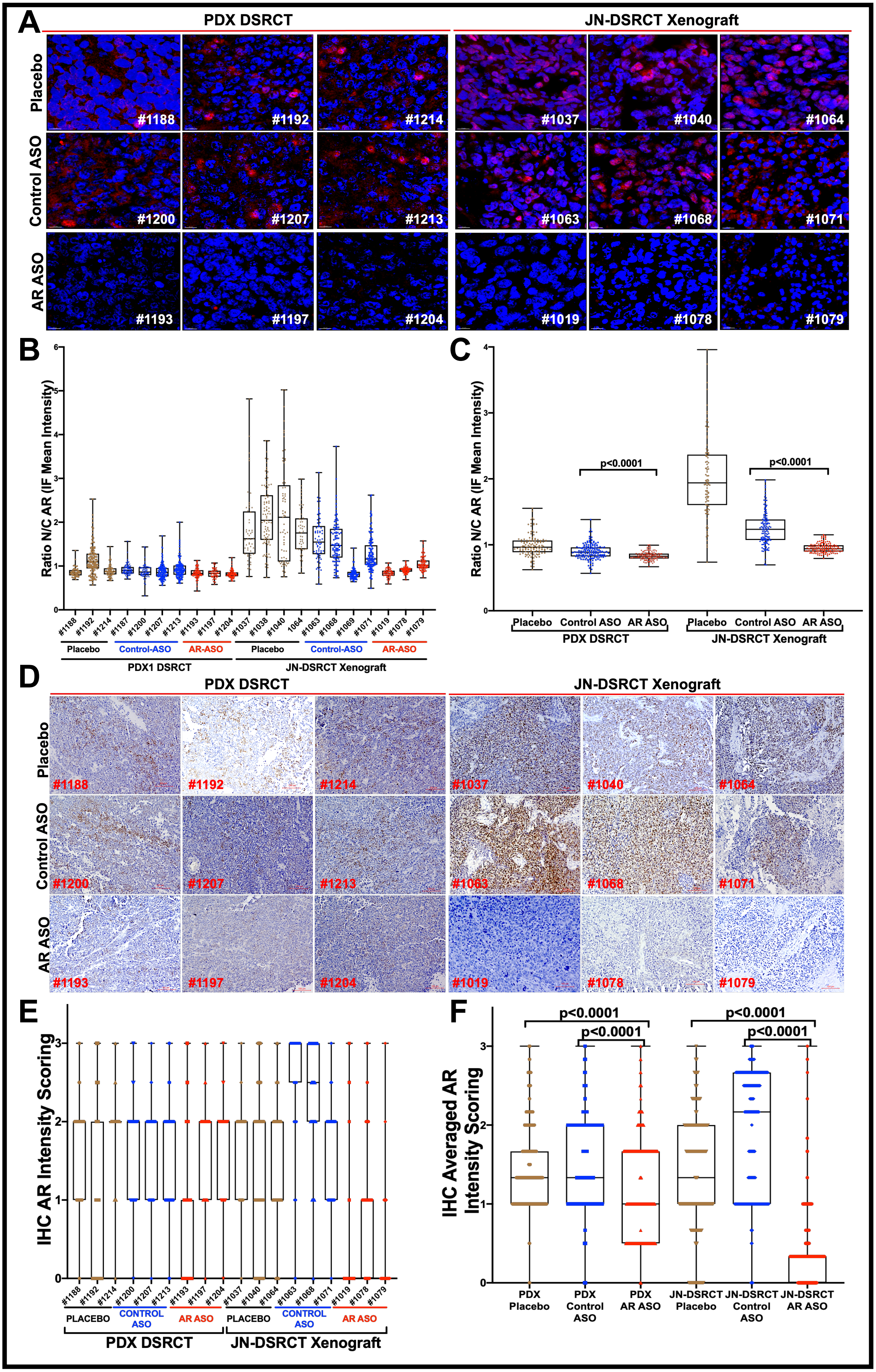

### Supplemental Figure 6

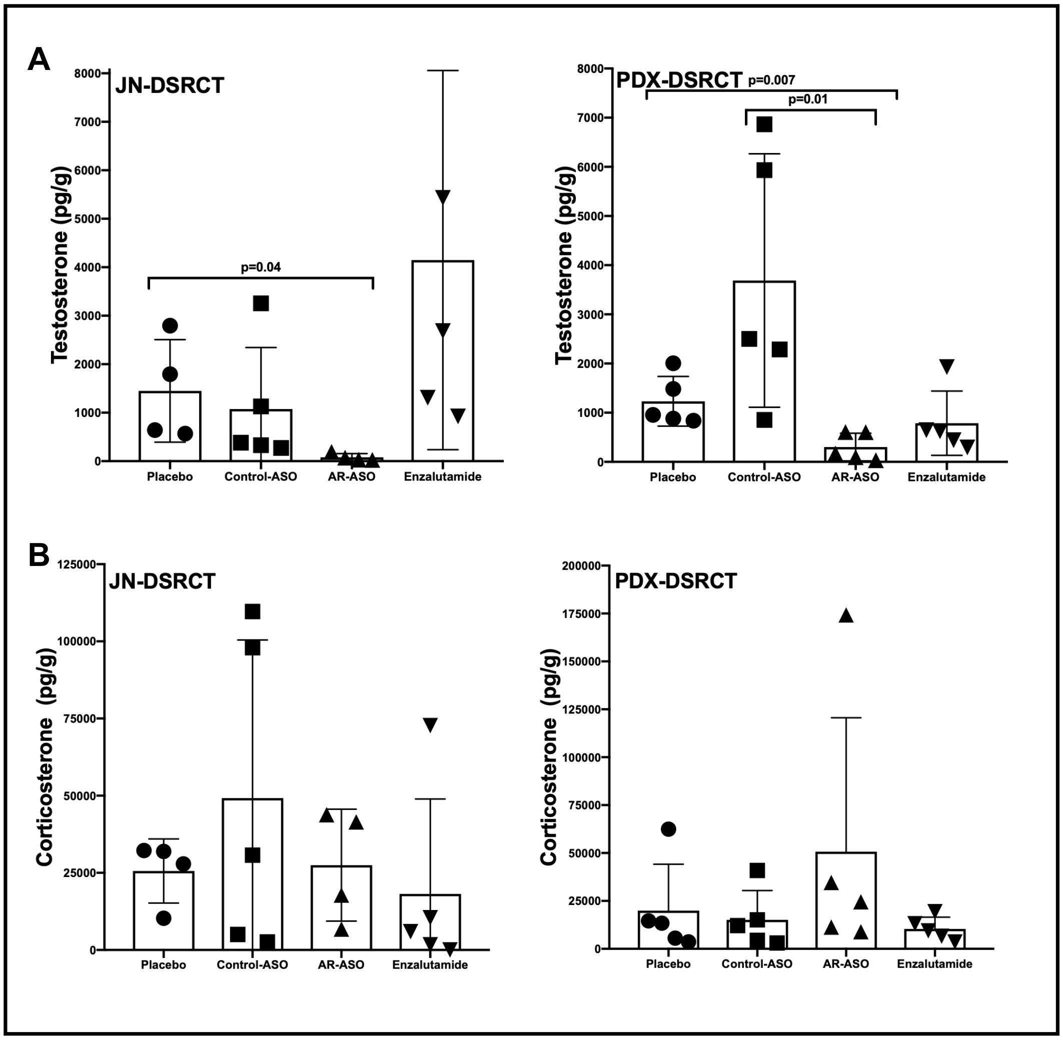

### Supplemental Figure 7

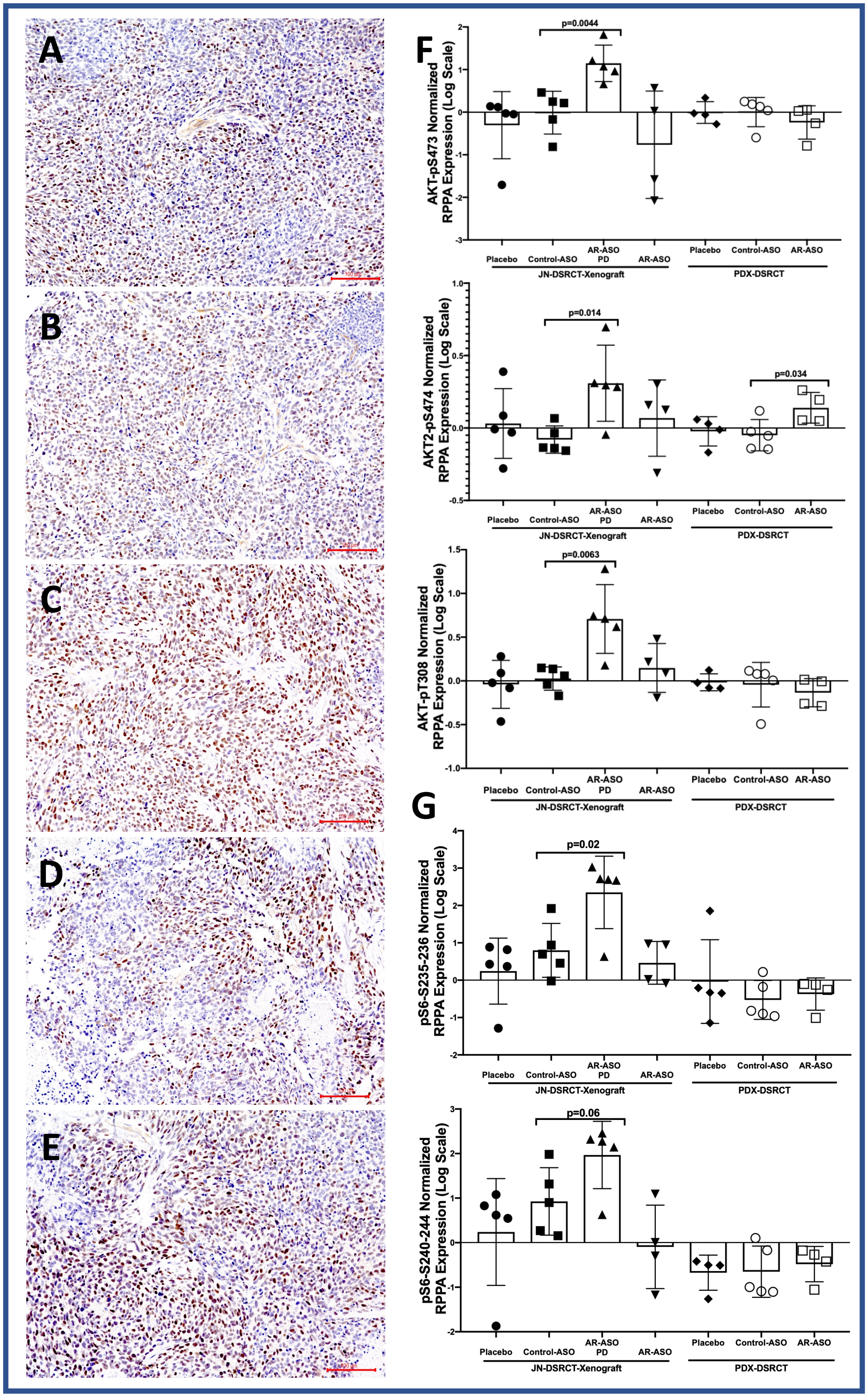

### Supplemental Figure 8

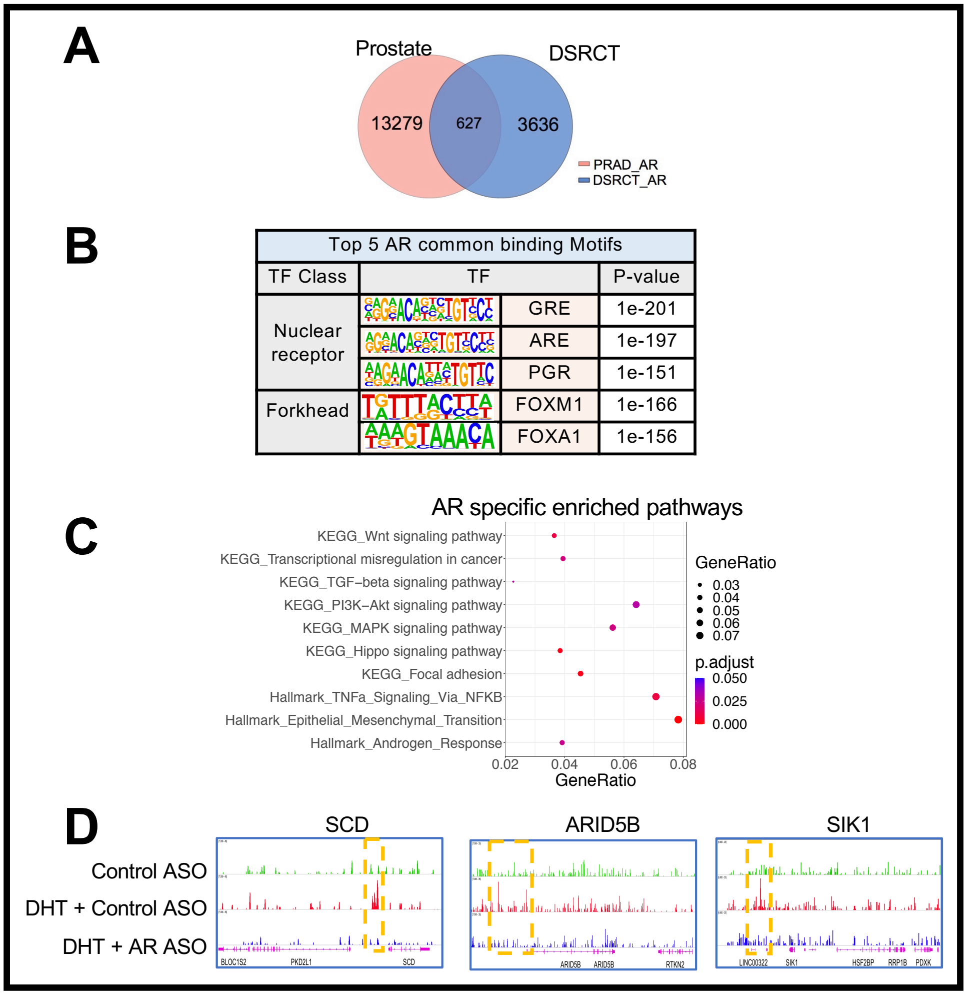

### Supplementary Figure 9

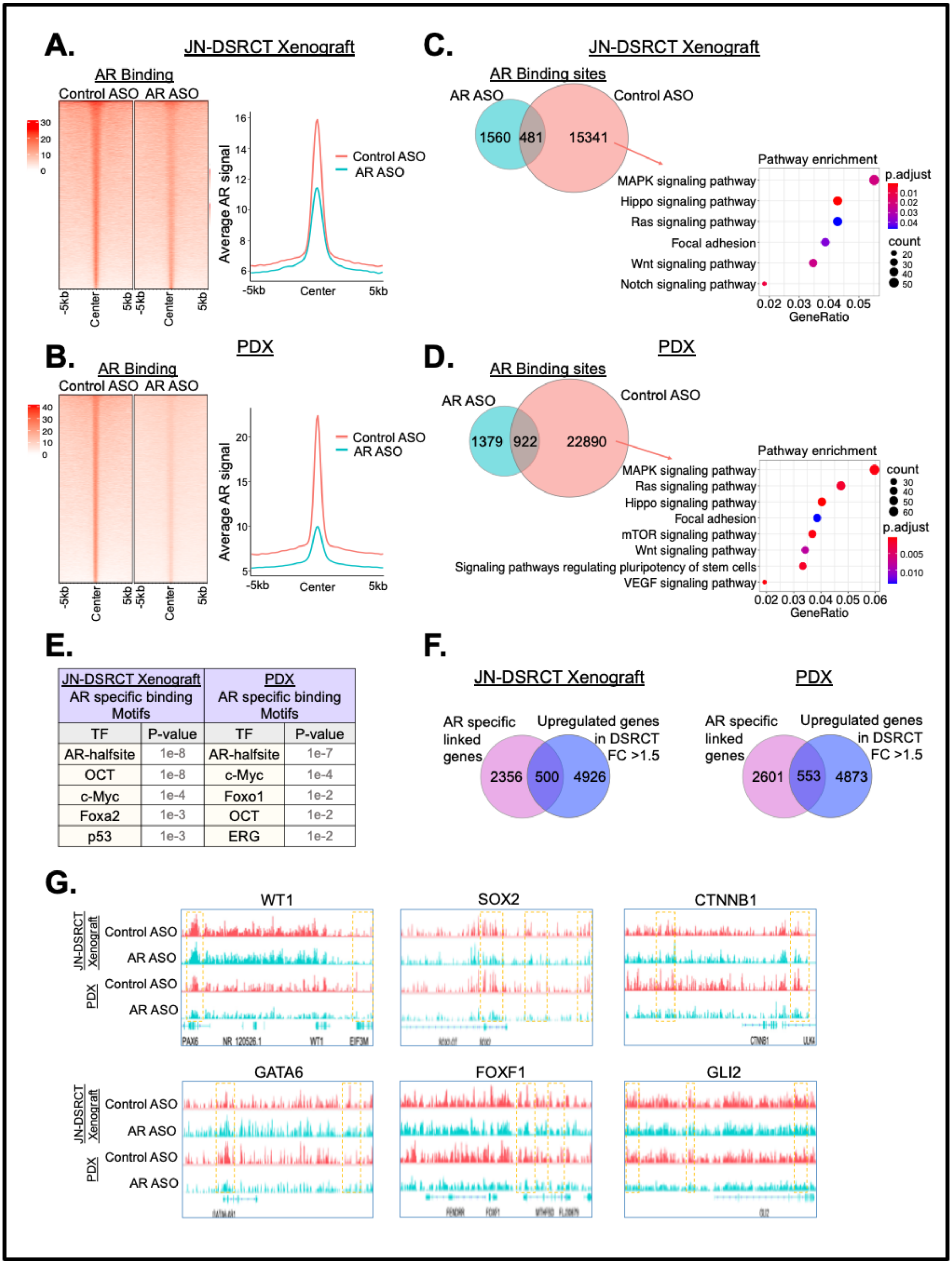


| JN-DSRCT Xenograft AR specific binding Motifs | | PDX AR specific binding Motifs | |
|---|---|---|---|
| TF | P-value | TF | P-value |
| AR-halfsite | 1e-8 | AR-halfsite | 1e-7 |
| OCT | 1e-8 | c-Myc | 1e-4 |
| c-Myc | 1e-4 | Foxo1 | 1e-2 |
| Foxa2 | 1e-3 | OCT | 1e-2 |
| p53 | 1e-3 | ERG | 1e-2 |

### Supplemental Figure 10

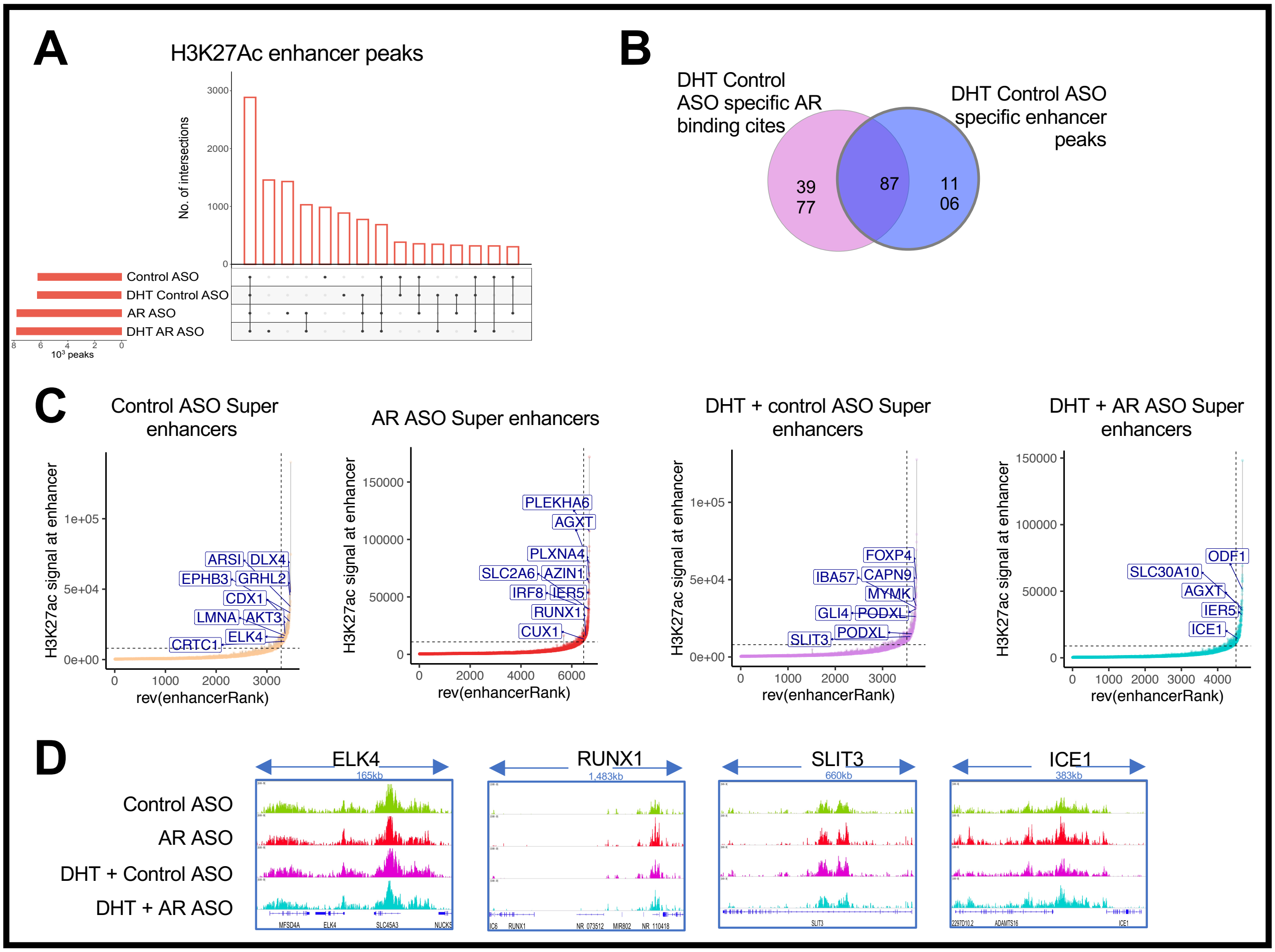

### Supplementary Figure 11

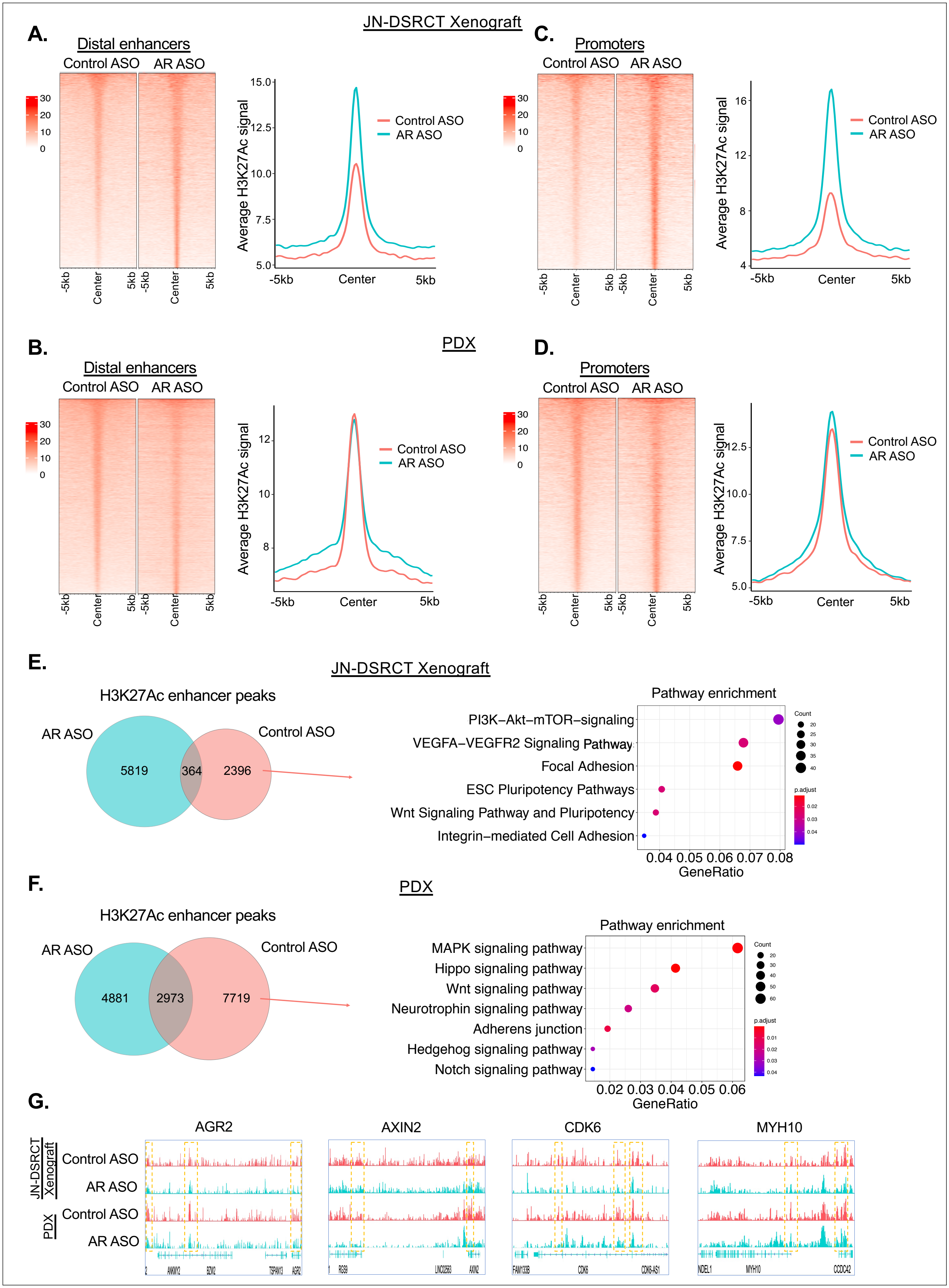

#### Supplemental Figure 12

A

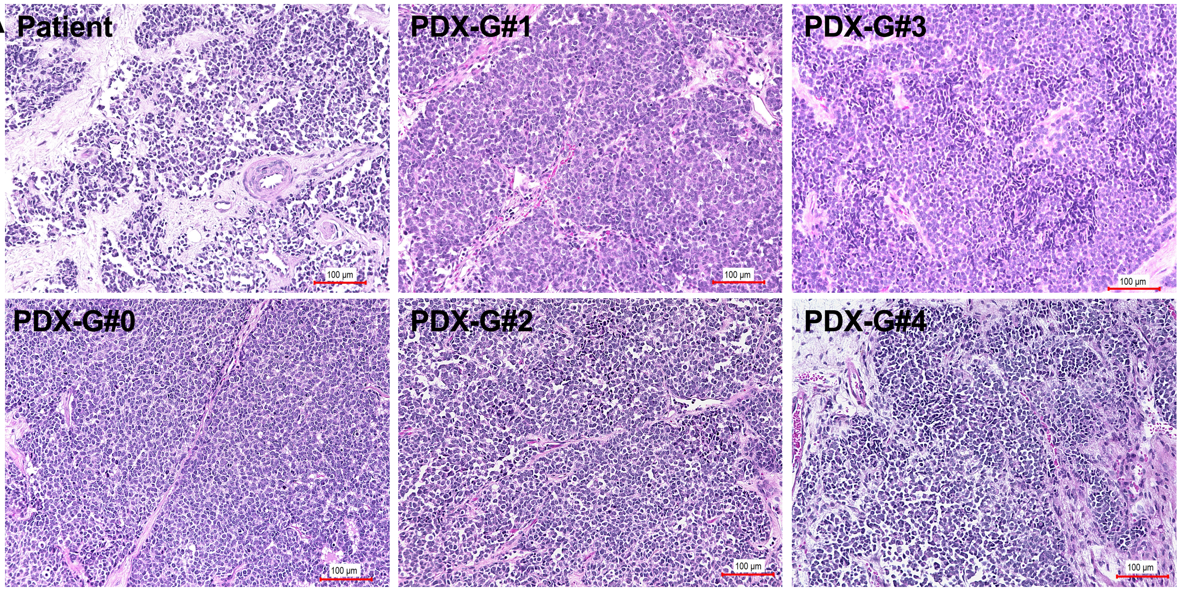


B

| | AMEL | CSF1PO | D13S317 | D16S539 | D18S51 | D21S11 | D3S1358 | D5S818 | D7S820 | D8S1179 | FGA | TH01 | TPOX | vWA |
|---|---|---|---|---|---|---|---|---|---|---|---|---|---|---|
| **Patient** | X,Y | 10,12 | 8 | 10,13 | 12,14 | 30 | 14 | 13 | 9 | 11,12 | 25,26 | 6,7 | 8,9 | 17 |
| **PDX-G0** | X,Y | 10,12 | 8 | 10,13 | 12,14 | 30 | 14 | 13 | 9 | 11,12 | 25,26 | 7 | 8,9 | 17 |
| **PDX-G1** | X,Y | 10,12 | 8 | 10,13 | 12,14 | 30 | 14 | 13 | 9 | 11,12 | 25,26 | 7 | 8,9 | 17 |
| **PDX-G2** | X,Y | 10,12 | 8 | 10,13 | 12,14 | 30 | 14 | 13 | 9 | 11,12 | 25,26 | 7 | 8,9 | 17 |
| **PDX-G3** | X,Y | 10,12 | 8 | 10,13 | 12,14 | 30 | 14 | 13 | 9 | 11,12 | 25,26 | 7 | 8,9 | 17 |
| **PDX-G4** | X,Y | 10,12 | 8 | 10,13 | 12,14 | 30 | 14 | 13 | 9 | 11,12 | 25,26 | 7 | 8,9 | 17 |

C

| Sample_ID | Gender | DOB | Diagnosis Date | Age @ Diagnosis | Race | Pathology Type | Pathology Grade | Metastatic Disease @ Diagnosis | Metastatic Site | Treatment Modalities | Vital Status |
|---|---|---|---|---|---|---|---|---|---|---|---|
| **AHJ-T15** | **Male** | **1/1/04** | **9/2/13** | **9.7** | **White** | **Metastatic** | **High** | **Yes** | **Lung** | **Chemotherapy Surgery** | **Unknown** |

#### SUPPLEMENTARY MATERIALS AND METHODS

##### Antibodies used for Western blots

The antibodies used in western blots are listed below

| Antibodies | Company |
|---|---|
| AR, SYK, TAU, c-Kit, pS6-S235/236, b-Actin, GAPDH, IGFBP2, AMPK-pT172, AKT, Tuberin, ITGAV, ITGA4, ITGA5, ITGB5, ITGB3, ITGB1, NCOA1, NCOA2, NCOA3 | Cell Signaling Technologies (CST) |

##### Antibodies used for RPPA Analyses

| # | Official Ab Name | Ab Name Reported on Dataset | Gene Name | Company |
|---|---|---|---|---|
| 1 | 14-3-3 beta | 14-3-3-beta | YWHAB | Santa Cruz |
| 2 | 14-3-3 epsilon | 14-3-3-epsilon | YWHAE | Santa Cruz |
| 3 | 14-3-3 zeta | 14-3-3-zeta | YWHAZ | Santa Cruz |
| 4 | 4E-BP1 | 4E-BP1 | EIF4EBP1 | CST |
| 5 | 4E-BP1_pS65 | 4E-BP1_pS65 | EIF4EBP1 | CST |
| 6 | 4E-BP1_pT37_T46 | 4E-BP1_pT37_T46 | EIF4EBP1 | CST |
| 7 | 53BP1 | 53BP1 | TP53BP1 | CST |
| 8 | Acetyl CoA Carboxylase 1 | ACC1 | ACACA | Epitomics |
| 9 | Acetyl-CoA Carboxylase_pS79 | ACC_pS79 | ACACA ACACB | CST |
| 10 | ACVRL1 | ACVRL1 | ACVRL1 | Abcam |
| 11 | ADAR1 | ADAR1 | ADAR | Abcam |
| 12 | Akt | Akt | AKT1 AKT2 AKT3 | CST |
| 13 | Akt_pS473 | Akt_pS473 | AKT1 AKT2 AKT3 | CST |
| 14 | Akt_pT308 | Akt_pT308 | AKT1 AKT2 AKT3 | CST |
| 15 | AMPKa | AMPK-alpha | PRKAA1 | CST |
| 16 | AMPKa_pT172 | AMPK-alpha_pT172 | PRKAA1 | CST |
| 17 | Androgen Receptor | AR | AR | Epitomics |
| 18 | Annexin I | Annexin-I | ANXA1 | BD Biosciences |
| 19 | Annexin VII | Annexin-VII | ANXA7 | BD Biosciences |
| 20 | A-Raf | A-Raf | ARAF | CST |
| 21 | ARHI | ARHI | DIRAS3 | MDACC Laboratory |
| 22 | ATM | ATM | ATM | CST |
| 23 | ATM_pS1981 | ATM_pS1981 | ATM | CST |
| 24 | ATP5H | ATP5H | ATP5H | Abcam |
| 25 | ATR | ATR | ATR | CST |
| 27 | Bak | Bak | BAK1 | Epitomics |
| 28 | BAP1 | BAP1 | BAP1 | Santa Cruz |
| 29 | Bax | Bax | BAX | CST |
| 30 | BCL2 | Bcl2 | BCL2 | Dako |
| 31 | Bcl-xL | Bcl-xL | BCL2L1 | CST |
| 32 | Beclin | Beclin | BECN1 | Santa Cruz |

| # | Official Ab Name | Ab Name Reported on Dataset | Gene Name | Company |
|---|---|---|---|---|
| 33 | beta-Catenin | b-Catenin | CTNNB1 | CST |
| 34 | beta-Catenin_pT41_S45 | b-Catenin_pT41_S45 | CTNNB1 | CST |
| 35 | Bid | Bid | BID | Abcam |
| 36 | Bim | Bim | BCL2L11 | Abcam |
| 37 | B-Raf | B-Raf | BRAF | Santa Cruz |
| 38 | B-Raf_pS445 | B-Raf_pS445 | BRAF | CST |
| 39 | BRCA2 | BRCA2 | BRCA2 | CST |
| 40 | Caspase-7 cleavedD198 | Caspase-7-cleaved | CASP7 | CST |
| 41 | Caspase-8 | Caspase-8 | CASP8 | CST |
| 42 | Caveolin-1 | Caveolin-1 | CAV1 | CST |
| 43 | CD29 | CD29 | ITGB1 | BD Biosciences |
| 44 | CD31 | CD31 | PECAM1 | Dako |
| 45 | CD49b | CD49b | ITGA2 | BD Biosciences |
| 46 | cdc2/CDK1 | CDK1 | CDC2-<br>CDK1 | CST |
| 47 | CDKN2A/p16INK4a | p16INK4a | CDKN2A | Abcam |
| 48 | Chk1 | Chk1 | CHEK1 | CST |
| 49 | Chk1_pS345 | Chk1_pS345 | CHEK1 | CST |
| 50 | Chk2 | Chk2 | CHEK2 | CST |
| 51 | Chk2_pT68 | Chk2_pT68 | CHEK2 | CST |
| 52 | c-Jun_pS73 | c-Jun_pS73 | JUN | CST |
| 53 | c-Kit | c-Kit | KIT | Abcam |
| 54 | Claudin 7 | Claudin-7 | CLDN7 | Novus Biologicals |
| 55 | c-Met | c-Met | MET | CST |
| 56 | c-Met_pY1234_Y1235 | c-Met_pY1234_Y1235 | MET | CST |
| 57 | c-Myc | c-Myc | MYC | Santa Cruz |
| 58 | COL6A1 | Collagen-VI | COL6A1 | Santa Cruz |
| 59 | Complex II Subunit | Complex-II-Subunit | SDHA | Invitrogen |
| 60 | Cox IV | Cox-IV | COX4I1 | Abcam |
| 61 | Cox2 | Cox2 | PTGS2 | CST |
| 62 | C-Raf/Raf-1 | C-Raf | RAF1 | Millipore |
| 63 | C-Raf_pS338 | C-Raf_pS338 | RAF1 | CST |
| 64 | Cyclin B1 | Cyclin-B1 | CCNB1 | Epitomics |
| 65 | Cyclin D1 | Cyclin-D1 | CCND1 | Santa Cruz |
| 66 | Cyclin E1 | Cyclin-E1 | CCNE1 | Santa Cruz |
| 67 | Cyclophilin F | Cyclophilin-F | PPIF | Abcam |
| 68 | Dvl3 | Dvl3 | DVL3 | CST |
| 69 | E2F-1 | E2F1 | E2F1 | Santa Cruz |
| 70 | E-Cadherin | E-Cadherin | CDH1 | CST |
| 71 | eEF2 | eEF2 | EEF2 | CST |
| 72 | eEF2K | eEF2K | EEF2K | CST |
| 73 | EGFR | EGFR | EGFR | CST |
| 74 | EGFR_pY1068 | EGFR_pY1068 | EGFR | CST |
| 75 | EGFR_pY1173 | EGFR_pY1173 | EGFR | Abcam |
| 76 | eIF4E | eIF4E | EIF4E | CST |
| 77 | eIF4G | eIF4G | EIF4G1 | CST |
| 78 | ER alpha_pS118 | ER-alpha_pS118 | ESR1 | Epitomics |
| 79 | ErbB2/HER2 | HER2 | ERBB2 | Lab Vision |

| # | Official Ab Name | Ab Name Reported on Dataset | Gene Name | Company |
|---|---|---|---|---|
| 80 | ErbB2/HER2_pY1248 | HER2_pY1248 | ERBB2 | R&D Systems |
| 81 | ErbB3/HER3 | HER3 | ERBB3 | Santa Cruz |
| 82 | ErbB3/HER3_pY1289 | HER3_pY1289 | ERBB3 | CST |
| 83 | ERCC1 | ERCC1 | ERCC1 | Santa Cruz |
| 84 | ERRFI1/MIG6 | MIG6 | ERRFI1 | Sigma-Aldrich |
| 85 | Estrogen Receptor | ER-alpha | ESR1 | Lab Vision |
| 86 | Ets-1 | Ets-1 | ETS1 | Bethyl |
| 87 | FAK | FAK | PTK2 | CST |
| 88 | FAK_pY397 | FAK_pY397 | PTK2 | CST |
| 89 | Fatty Acid Synthase | FASN | FASN | CST |
| 90 | Fibronectin | Fibronectin | FN1 | Epitomics |
| 91 | FoxM1 | FoxM1 | FOXM1 | CST |
| 92 | FoxO3a | FoxO3a | FOXO3 | CST |
| 93 | FoxO3a_pS318_S321 | FoxO3a_pS318_S321 | FOXO3 | CST |
| 94 | G6PD | G6PD | G6PD | Santa Cruz |
| 95 | Gab2 | Gab2 | GAB2 | CST |
| 96 | GAPDH | GAPDH | GAPDH | Life Technologies |
| 97 | GATA3 | GATA3 | GATA3 | BD Biosciences |
| 98 | GCN5L2 | GCN5L2 | KAT2A | CST |
| 99 | Glycogen Synthase | Gys | GYS1 | CST |
| 100 | Glycogen Synthase_pS641 | Gys_pS641 | GYS1 | CST |
| 101 | GPBB | GPBB | PYGB | Novus Biologicals |
| 102 | GSK-3alpha/beta | GSK-3ab | GSK3A GSK3B | Santa Cruz |
| 103 | GSK-3alpha/beta_pS21_S9 | GSK-3ab_pS21_S9 | GSK3A GSK3B | CST |
| 104 | GSK-3beta_pS9 | GSK-3b_pS9 | GSK3B | CST |
| 105 | Heregulin | Heregulin | NRG1 | CST |
| 106 | HIAP | HIAP | BIRC2 | Millipore |
| 107 | Histone H3 | Histone-H3 | H3F3A H3F3B | Abcam |
| 108 | IGF-1Receptor beta | IGF1R-beta | IGF1R | CST |
| 109 | IGFBP2 | IGFBP2 | IGFBP2 | CST |
| 110 | INPP4b | INPP4b | INPP4B | CST |
| 111 | IRS1 | IRS1 | IRS1 | Millipore |
| 112 | JAB1 | JAB1 | COPS5 | Santa Cruz |
| 113 | JNK/SAPK_pT183_Y185 | JNK_pT183_Y185 | MAPK8 | CST |
| 114 | JNK2 | JNK2 | MAPK9 | CST |
| 115 | KMT3A/HYPB/HIF 1 | SETD2 | SETD2 | Abcam |
| 116 | Lck | Lck | LCK | CST |
| 117 | MAPK_pT202_Y204 | MAPK_pT202_Y204 | MAPK1 MAPK3 | CST |
| 118 | Mcl 1 | Mcl-1 | MCL1 | CST |
| 119 | MDM2_pS166 | MDM2_pS166 | MDM2 | CST |
| 120 | MEK1 | MEK1 | MAP2K1 | Epitomics |
| 121 | MEK1_pS217_S221 | MEK1_pS217_S221 | MAP2K1 MAP2K2 | CST |

| # | Official Ab Name | Ab Name Reported on Dataset | Gene Name | Company |
|---|---|---|---|---|
| 122 | MEK2 | MEK2 | MAP2K2 | CST |
| 123 | Merlin/NF2 | Merlin | NF2 | Novus Biologicals |
| 124 | MSH2 | MSH2 | MSH2 | CST |
| 125 | MSH6 | MSH6 | MSH6 | Novus Biologicals |
| 126 | mTOR | mTOR | MTOR | CST |
| 127 | mTOR_pS2448 | mTOR_pS2448 | MTOR | CST |
| 128 | Myosin heavy chain 11 | Myosin-11 | MYH11 | Novus Biologicals |
| 129 | Myosin IIa_pS1943 | Myosin-IIa_pS1943 | MYH9 | CST |
| 130 | NAPSIN A | NAPSIN-A | NAPSA | Abcam |
| 131 | N-Cadherin | N-Cadherin | CDH2 | CST |
| 132 | NDRG1_pT346 | NDRG1_pT346 | NDRG1 | CST |
| 133 | NF-kappaB p65_pS536 | NF-kB-p65_pS536 | RELA | CST |
| 134 | Notch1 | Notch1 | NOTCH1 | CST |
| 135 | N-Ras | N-Ras | NRAS | Santa Cruz |
| 136 | p21 | p21 | CDKN1A | Santa Cruz |
| 137 | p27 KIP 1 | p27-Kip-1 | CDKN1B | Abcam |
| 138 | p27 KIP 1_pT198 | p27_pT198 | CDKN1B | Abcam |
| 139 | p27/Kip1_pT157 | p27_pT157 | CDKN1B | R&D Systems |
| 140 | p38 alpha MAPK | p38-alpha | MAPK14 | CST |
| 141 | p38 MAPK | p38 | MAPK14 | CST |
| 142 | p38 MAPK_pT180_Y182 | p38_pT180_Y182 | MAPK14 | CST |
| 143 | p53 | p53 | TP53 | CST |
| 144 | p70 S6 Kinase_pT389 | p70-S6K_pT389 | RPS6KB1 | CST |
| 145 | p70/S6K1 | p70-S6K1 | RPS6KB1 | Epitomics |
| 146 | PAI-1 | PAI-1 | SERPINE1 | BD Biosciences |
| 147 | PARK7/DJ1 | DJ1 | PARK7 | Abcam |
| 148 | PARP cleavedD214 | PARP-cleaved | PARP1 | CST |
| 149 | PARP-1 | PARP1 | PARP1 | Santa Cruz |
| 150 | Paxillin | Paxillin | PXN | Epitomics |
| 151 | PCNA | PCNA | PCNA | Abcam |
| 152 | Pdcd-1L1 | Pdcd-1L1 | CD274 | Santa Cruz |
| 153 | Pdcd4 | Pdcd4 | PDCD4 | Rockland |
| 154 | PDGFR beta | PDGFR-beta | PDGFRB | CST |
| 155 | PDK1 | PDK1 | PDPK1 | CST |
| 156 | PDK1_pS241 | PDK1_pS241 | PDPK1 | CST |
| 157 | PEA-15 | PEA-15 | PEA15 | CST |
| 158 | PED/PEA-15_pS116 | PEA-15_pS116 | PEA15 | Invitrogen |
| 159 | PI3 Kinase p110 alpha | PI3K-p110-alpha | PIK3CA | CST |
| 160 | PI3K p85 | PI3K-p85 | PIK3R1 | Millipore |
| 161 | PKC alpha | PKC-alpha | PRKCA | Millipore |
| 162 | PKC alpha_pS657 | PKC-alpha_pS657 | PRKCA | Millipore |
| 163 | PKC beta II_pS660 | PKC-beta-II_pS660 | PRKCA<br>PRKCB<br>PRKCD<br>PRKCE<br>PRKCH<br>PRKCQ | CST |

| # | Official Ab Name | Ab Name Reported on Dataset | Gene Name | Company |
|---|---|---|---|---|
| 164 | PKC delta_pS664 | PKC-delta_pS664 | PRKCD | Millipore |
| 165 | PMS2 | PMS2 | PMS2 | Novus Biologicals |
| 166 | PRAS40_pT246 | PRAS40_pT246 | AKT1S1 | Life Technologies |
| 167 | PREX1 | PREX1 | PREX1 | Abcam |
| 168 | Progesterone Repector | PR | PGR | Abcam |
| 169 | PTEN | PTEN | PTEN | CST |
| 170 | Rab11 | Rab11 | RAB11A<br>RAB11B | CST |
| 171 | Rab25 | Rab25 | RAB25 | CST |
| 172 | Rad50 | Rad50 | RAD50 | Millipore |
| 173 | Rad51 | Rad51 | RAD51 | CST |
| 174 | Raptor | Raptor | RPTOR | CST |
| 175 | Rb | Rb | RB1 | CST |
| 176 | Rb_pS807_S811 | Rb_pS807_S811 | RB1 | CST |
| 177 | RBM15 | RBM15 | RBM15 | Novus Biologicals |
| 178 | Rictor | Rictor | RICTOR | CST |
| 179 | Rictor_pT1135 | Rictor_pT1135 | RICTOR | CST |
| 180 | RSK | RSK | RPS6KA1<br>RPS6KA2<br>RPS6KA3 | CST |
| 181 | S6_pS235_S236 | S6_pS235_S236 | RPS6 | CST |
| 182 | S6_pS240_S244 | S6_pS240_S244 | RPS6 | CST |
| 183 | SCD | SCD | SCD | Santa Cruz |
| 184 | SF2/ASF | SF2 | SRSF1 | Invitrogen |
| 185 | Shc_pY317 | Shc_pY317 | SHC1 | CST |
| 186 | Smac/Diablo | Smac | DIABLO | CST |
| 187 | Smad1 | Smad1 | SMAD1 | Epitomics |
| 188 | Smad3 | Smad3 | SMAD3 | Abcam |
| 189 | Smad4 | Smad4 | SMAD4 | Santa Cruz |
| 190 | Snail | Snail | SNAI1 | CST |
| 191 | Src | Src | SRC | Millipore |
| 192 | Src Family_pY416 | Src_pY416 | SRC<br>LYN<br>FYN<br>LCK<br>YES1<br>HCK | CST |
| 193 | Src_pY527 | Src_pY527 | SRC<br>YES1<br>FYN<br>FGR | CST |
| 194 | Stat3_pY705 | Stat3_pY705 | STAT3 | CST |
| 195 | Stat5a | Stat5a | STAT5A | Abcam |
| 196 | Stathmin 1 | Stathmin-1 | STMN1 | Abcam |
| 197 | Syk | Syk | SYK | Santa Cruz |
| 198 | TAZ | TAZ | WWTR1 | CST |
| 199 | TIGAR | TIGAR | C12ORF5 | Abcam |
| 200 | Transferrin Receptor | TFRC | TFRC | Novus Biologicals |

| # | Official Ab Name | Ab Name Reported on Dataset | Gene Name | Company |
|---|---|---|---|---|
| 201 | Transglutaminase II | Transglutaminase | TGM2 | Lab Vision |
| 202 | TSC1/Hamartin | TSC1 | TSC1 | CST |
| 203 | TSC2 Tuberin_pT1462 | Tuberin_pT1462 | TSC2 | CST |
| 204 | TTF1 | TTF1 | NKX2-1 | Abcam |
| 205 | Tuberin | Tuberin | TSC2 | Epitomics |
| 206 | Twist | TWIST | TWIST2 | Santa Cruz |
| 207 | Tyro3 | Tyro3 | TYRO3 | CST |
| 208 | UBAC1 | UBAC1 | UBAC1 | Sigma-Aldrich |
| 209 | UGT1A | UGT1A | UGT1A1 | Santa Cruz |
| 210 | UQCRC2 | UQCRC2 | UQCRC2 | Abcam |
| 211 | VDAC1/Porin | Porin | VDAC1 | Abcam |
| 212 | VEGF Receptor 2 | VEGFR-2 | KDR | CST |
| 213 | VHL | VHL | VHL | BD Biosciences |
| 214 | XRCC1 | XRCC1 | XRCC1 | CST |
| 215 | YAP | YAP | YAP1 | Santa Cruz |
| 216 | YAP_pS127 | YAP_pS127 | YAP1 | CST |
| 217 | YB1 | YB1 | YBX1 | Novus Biologicals |
| 218 | YB1_pS102 | YB1_pS102 | YBX1 | CST |

**RPPA and western blot Analyses**

Raw log2 intensity values were normalized for global protein expression by median centering across 151 antibodies tested. Individual protein expressions in DSRCT specimens, ES specimens, and mesenteric normal tissue specimens were compared using two-sided unpaired t-tests with the GeneSpring GX software program version 12.6.1-GX (Agilent Technologies). For multiple comparison testing, proteins whose expression levels between specimen types were 2-fold and significantly different ($p \leq 0.05$), were subjected to unsupervised hierarchical clustering. Differentially expressed proteins identified by RPPA were confirmed by Western blot analysis. The RPPA data for the proteomic comparison of DSRCT patient tumors, normal mesenteric tissues, and Ewing sarcoma patient tumors are available from the Gene Expression Omnibus repository under GSE108687 series.

The preparation of extracted protein from cells, DSRCT patient tumors, or xenograft animal tumors for western blotting validations was prepared as described previously[1]. Lysis buffer (1% Triton X-100, 50 mM HEPES, pH 7.4, 150 mM NaCl, 1.5 mM $MgCl_2$, 1 mM EGTA, 100 mM NaF, 10 mM Na pyrophosphate, 1 mM $Na_3VO_4$, 10% glycerol) containing freshly added protease and phosphatase inhibitors (Roche Applied Sciences) was applied to lyse cellular washed pellets via cold incubation. Next, protein extraction from xenograft tumors was performed by homogenizing approximately 10 mg of frozen tissue in 500 µL of the lysis buffer using an electric tissue homogenizer (PRO Scientific). The homogenized tumors were incubated at +4°C for 2 hours to complete their dissociation and lysis. Altogether, the total lysed proteins from tumors were collected after centrifugation, quantified using BCA protein assay kit (Thermo Fisher Scientific), and stored at -80°C until further analyses. Western blots were performed as previously described by our group [1,2]. Protein detection was performed using a list of antibodies provided in the above supplementary table of antibodies used for Western blots. The immune-reactive proteins were captured using horseradish peroxidase-conjugated secondary anti-rabbit IgG or anti-mouse IgG antibodies (Cell Signaling Technology), amplified using the SuperSignal West Dura chemiluminescent substrate (Thermo Fisher Scientific), detected using the Chemi-Doc system (Bio-Rad), and quantified for their densitometry using the ImageJ Gel Analysis tool (NIH, Bethesda, MD).

**TMA Preparation and Immunohistochemistry Analyses**

Four-micrometer unstained slides were prepared from this tissue microarray. Immunohistochemical studies were performed using an autostainer (Bond-Max; Leica Microsystems, Buffalo Grove, IL, USA) with anti-AR (1:30; clone AR441, Dako#M3562) antibody. This primary antibody was detected using a Bond polymer refine detection kit according to the manufacturer protocol (Leica, #DS9800). Then, the slides were dehydrated in grade alcohols, cleared in xylene, and cover-slipped. The TMA staining results were scored on intensity (low, moderate, and high) as well as the percentage of tumoral labeling (Positive (>50%), Low positive (10-50%), Focal (1-10%), and Negative (0-1%)).

**WST1 Cell Proliferation Assays**

The cells cultivated for 24 hours to let them adhere, before adding serially diluted concentrations of dihydrotestosterone (DHT, 0.0064-2000nM; Selleckchem), or AR-based drug treatments (Enzalutamide, AR antisense oligonucleotides; 0.032-5mM) for up to 48 hours or 72 hours, respectively. WST-1 added, and cells incubated for an additional 2 hours. Cell proliferation was measured at 450 nm in a microplate reader (DTX880, Beckman Coulter). In both assays, the agonistic effect of DHT or cytotoxic effect of the drugs was expressed as a percentage of cell proliferation. IC50 values were calculated by the sigmoidal dose-response curve-fit using Prism GraphPad 8.0.

**Immunohistochemistry and digital image analyses**

Primary tumors from the DSRCT xenograft and PDX1 group of mice, untreated and treated with control-ASO or AR-ASO, were fixed in 10% formalin, embedded in paraffin, and then sliced in 5µm sections. The EZ-retriever microwave-based pretreatment and antigen retrieval system (Biogenex, CA) used for dewaxing, rehydration, and antigen retrieval of these FFPE lung tissue sections. AR protein expression by IHC was evaluated on Leica Bond MAX Auto-stainer by using primary antibody, rabbit antibody anti-human AR (Cell Signaling Technologies, #5153). The primary antibody was detected according to the manufacturer protocol (Leica, #DS9800). Then, the slides were dehydrated in grade alcohols, cleared in xylene, cover-slipped, and imaged with Keyence Microscope (Keyence, Tokyo, Japan) at 20X resolution. The digital images were processed and quantified using the Visiopharm software version (2020.04) (Hoersholm, Denmark). An APP (Analysis Protocol Package) was designed to quantify cell-based DAB staining in DSRCT preclinical samples using traditional thresholding methods. The algorithm was built around a three-step approach 1) Pre-Processing: HDAB-DAB and HDAB-Hematoxylin features were used to detect AR Positive cells (Green masking) and Negative cells (Blue masking) respectively by setting pixel values. A median filter of size 5X5 was used for the proper segmentation of cells. 2) Post Processing: Additional steps were designed to enhance the performance of the APP. Change by shape excludes the artifacts; merged cells were separated by separate labels step, and certain clear areas in the cells were filled with fill holes step to fully mask the cells. 3) Output Variable: Mean intensity of each cell and from each sample was extracted and exported into a spreadsheet. Finally, the data were plotted on a Scatter Plot using GraphPad Prism software, version 8 showing the AR Mean Intensity of each cell from all the analyzed samples, and the AR Average mean intensity from each preclinical group of treated mice.

#### SUPPLEMENTAL FIGURE LEGENDS

**Supplemental Figure 1. Proteomic signature differentiates DSRCT from normal mesenteric tissues. A)** Proteomic profiling between DSRCT and normal mesenteric tissue specimens. The protein lysates from DSRCT (red) and normal mesenteric tissue (green) were subjected to RPPA analysis for 151 proteins and phosphoproteins (red, increased signal; green, decreased signal). Unsupervised double-hierarchical clustering using the Pearson correlation distance metric between proteins (rows) and Centroid linkage (a clustering method) separated the 16 samples into two groups (columns). The heat map of all 16 samples showed 20 proteins that distinguished DSRCT from normal mesenteric tissues ($p \leq 0.05$; fold-change ≥2). **B)** Western blotting was used to validate the proteins identified by RPPA as being differentially expressed between DSRCT and normal mesenteric tissue. **C)** Normalized protein expression is relative to $\beta$-actin.

**Supplemental Figure 2. Proteomic and genomic profiling of DSRCT and other types of sarcoma primary tumor samples. A-C)** Examples of TMA immunohistochemical labeling in desmoplastic small round cell tumor. **A)** A complete absence of tumoral nuclear labeling for AR (100x). **B)** Variable tumoral labeling for AR (200x), with a mixture of tumor cells positive (brown nuclei) and absent (blue nuclei) for nuclear labeling. The extent of labeling was estimated as the percentage of all labeled cells. **C)** Diffuse nuclear tumoral labeling for AR (200x). **D)** Heatmap of the KEGG Prostate Cancer pathway and **E)** the corresponding GSEA plot. **F)** Heatmap of the GO Androgen Receptor Signaling Pathway and **G)** the corresponding GSEA plot. The Z-score normalized expression levels of the genes that make up the KEGG Prostate Cancer and GO Androgen Receptor Signaling pathways were used in unsupervised clustering to generate the heatmaps. DSRCT samples are denoted in red while other sarcoma subtypes (WDLPS, SSLPS, MLPS, OS, and SS) are in gray.

**Supplemental Figure 3. Additional proteomic profiling of DSRCT samples using western blotting analyses. A)** Western blot analyses of integrin subclasses (ITGAV, ITGA4, ITGA5, ITGB1, ITGB3, and ITGB5) and GAPDH expression in 11 DSRCT cohort primary tumors. **B)** Western blot analyses of steroid receptor coactivators subclasses (NCOA1: SRC-1, NCOA2: SRC-2, and NCOA3: SRC-3) and GAPDH expression in 11 DSRCT cohort primary tumors. **C)** Western blot analyses of steroid receptor coactivators subclasses (NCOA1: SRC-1, NCOA2: SRC-2, and NCOA3: SRC-3) and GAPDH expression in JN-DSRCT and LNCaP PC cell lines.

**Supplemental Figure 4. DHT treatment of JN-DSRCT cells enhanced their sensitivity towards AR-based antisense blockade.** WST1 proliferation cell-based assay testing AR-ASO and control ASO sequences in a dose-dependent manner after 72 hours in the presence or the absence of DHT.

**Supplemental Figure 5. Immunofluorescence confocal and IHC evaluation of AR expression in JN-DSRCT and PDX tumors after AR-based antisense therapy. A)** Immunofluorescence evaluation images of preclinical JN-DSRCT and PDX tumor samples. **B)** Representative Nuclear/Cytoplasmic ratio AR immunofluorescence confocal microscopy quantification, within the single cell or, **C)** the averaged treated samples (placebo, control ASO, and AR ASO). Bars represent standard deviations. **D)** Immunohistochemical evaluation images of preclinical JN-DSRCT and PDX tumor samples. IHC stains for AR in primary tumors of JN-

DSRCT and PDX DSRCT mice after treatment with AR ASO, control ASO, and placebo. 100μm scale bars are shown. **E)** Representative IHC AR intensity scoring of the preclinical JN-DSRCT and PDX tumor samples, within the single cell or, **F)** the averaged treated samples (placebo, control ASO, and AR ASO). Bars represent standard deviations.

**Supplemental Fig 6. Testosterone and corticosterone quantification in DSRCT preclinical animal samples.** Testosterone (panel A) and corticosterone (panel B) levels (pg/g) in JN-DSCRT xenograft and patient-derived tumor explants (PDX-DSRCT). Testosterone levels were statistically dissimilar in JN-DSRCT and PDX-DSRCT placebo versus AR-ASO with P values of p=0.04 and p=0.007, respectively. In the PDX-DSRCT, the control-ASO and AR-ASO testosterone levels were statistically different (p=0.01). The corticosterone levels were similar across all treatments in both JN-DSRCT and PDX-DSRCT tumor models.

**Supplemental Fig 7. Proteomic evaluation of AR expression in preclinical JN-DSRCT tumors after enzalutamide-based therapy. A-E)** IHC stains for AR in primary tumors of JN-DSRCT mice after treatment with enzalutamide. Representative images from five enzalutamide-treated JN-DSRCT xenograft mice are shown. 100μm scale bars are shown. **F)** The mean expression intensity values of the three AKT-pS473, AKT2-pS474, and AKT-pT308 proteins associated with different group of mice treatments in JN-DSRCT xenograft and PDX animal models of DSRTC, and their statistical significance after normalization for global protein expression by median centering across 151 antibodies in the RPPA panel. **G)** The mean expression intensity values of the two pS6-S235-236, and pS6-S240-244 proteins associated with different group of mice treatments in JN-DSRCT xenograft and PDX animal models of DSRTC, and their statistical significance after normalization for global protein expression by median centering across 151 antibodies in the RPPA panel.

**Supplemental Figure 8. AR binding in DSRCT cells. A)** Venn diagram showing the overlap of AR peaks between DHT + Control ASO DSCRCT and PRAD 22RV1 samples to identify the shared AR binding sites. **B)** List of top 5 enriched transcription factor (TF) motifs in DSRCT and PRAD shared AR binding sites. Motifs are identified using HOMER. **C)** List of enriched pathways for genes that harbor AR binding sites in their vicinity in DHT-treated DSRCT cells. **D)** IGV images showing enrichment of AR peaks around SCD, ARID5B, and SIK1 genes using aggregate ChIP-seq profiles of Control ASO, DHT + Control ASO, and DHT + AR ASO samples.

**Supplemental Figure 9. AR binding in DSRCT Xenograft and PDX tumors. A-B)** Heatmaps (left panels) and average intensity curves (right panels) of ChIP-seq reads (RPKM; reads per kilobase of transcript per million mapped reads) for AR binding regions. AR binding sites are shown in a 10-kb window (centered on the middle of the binding site) in control ASO and AR ASO in DSRCT Xenograft (**A**) and PDX (**B**). **C-D)** Venn diagram (left) showing the overlap of all AR peaks between control ASO and AR ASO samples in DSRCT Xenograft (**C**) and PDX (**D**) to identify the AR-unique or shared binding sites. Dot plot (right) showing significantly enriched pathways for top 5000 AR specific binding sites in DSRCT Xenograft (**C**) and PDX (**D**). Dot size represents gene ratio, and colors represents adjusted p-values. **E)** List of top 5 enriched transcription factor (TF) motifs in AR specific binding sites in DSRCT Xenograft and PDX. Motifs are identified using HOMER. **F)** Venn diagram showing the overlap of annotated genes for AR specific gained enhancer peaks in DSRCT Xenograft (left) and PDX (right) and upregulated gene list for DSRCT tumors vs. other sarcoma tumors. **G)** IGV images showing enrichment of AR peaks around WT1, SOX2, CTNNB1, GATA6, FOXF1, and GLI2 genes using aggregate ChIP-seq profiles of Control ASO and AR-ASO samples in DSRCT Xenograft (top) and PDX (bottom).

**Supplemental Figure 10. Enhancer reprogramming by AR in JN-DSRCT cells. A)** Upset plots displaying unique and shared enhancer regions between Control ASO, AR ASO, DHT + Control ASO, and DHT + AR ASO samples. **B)** Venn diagram showing the overlap of AR specific enhancer peaks and AR peaks. **C)** Inflection plot indicating super-enhancers (SEs) identified in Control ASO, AR ASO, DHT + Control ASO, and DHT + AR ASO samples. **D)** IGV images showing enrichment of super-enhancer H3K27Ac peaks around ELK4, RUNX1, SLIT3, and ICE1 genes using aggregate ChIP-seq profiles of Control ASO, AR ASO, DHT + Control ASO and DHT + AR ASO samples.

**Supplemental Figure 11. Enhancer reprogramming by AR in DSRCT Xenograft and PDX tumors. A-B)** Heatmaps (left panels) and average intensity curves (right panels) of ChIP-seq reads (RPKM; reads per kilobase of transcript per million mapped reads) for distal enhancer regions. H3K27Ac binding sites are shown in a 10-kb window (centered on the middle of the binding site) in Control ASO and AR ASO in DSRCT Xenograft (**A**) and PDX (**B**). **C-D)** Heatmaps (left panels) and average intensity curves (right panels) of ChIP-seq reads (RPKM; reads per kilobase of transcript per million mapped reads) for promoter regions. H3K27Ac binding sites are shown in a 10-kb window (centered on the middle of the binding site) in Control ASO and AR ASO in DSRCT Xenograft (**C**) and PDX (**D**). **E-F)** Venn diagram (left) showing the overlap of all enhancer peaks between control ASO and AR ASO samples in DSRCT Xenograft (**E**) and PDX (**F**) to identify the H3K27Ac-unique or shared binding sites. Dot plot (right) showing significantly enriched pathways for top 5000 H3K27Ac specific binding sites in DSRCT Xenograft (**E**) and PDX (**F**). Dot size represents gene ratio, and colors represents adjusted p-values. **G**) IGV images showing enrichment of H3K27Ac peaks around AGR2, AXIN2, CDK6, and MYH10 genes using aggregate ChIP-seq profiles of Control ASO and AR-ASO samples in DSRCT Xenograft (top) and PDX (bottom).

**Supplemental Figure 12. Histologic and genetic analyses of DSRCT patient and PDX tumors. A)** The DSRCT patient and his $4^{th}$-generation PDX demonstrated similar histological features by H&E staining. **B)** STR profiling of the PDX matched the patient. **C)** Clinicopathological characteristics of the human tumor used for DSRCT PDX generation.

**SUPPLEMENTAL TABLES**

**Supplemental Table 1.** AR specific binding sites and motif enrichment in JN-DSRCT cells.

**Supplemental Table 2.** AR-dependent enhancer reprogramming in JN-DSRCT cells.

**Supplemental Table 3.** Super enhancers and regulated genes in JN-DSRCT cells.

**Supplemental Table 4.** Demographic information for DSRCT and Ewing sarcoma patients in RPPA analysis.

**Supplemental Table 5.** Normalized RPPA data between DSRCT and paired adjacent normal-appearing mesenteric tissues from the same patients
